# Piximi - An Images to Discovery web tool for bioimages and beyond

**DOI:** 10.1101/2024.06.03.597232

**Authors:** Levin M Moser, Nodar Gogoberidze, Andréa Papaleo, Alice Lucas, David Dao, Christoph A Friedrich, Lassi Paavolainen, Csaba Molnar, David R Stirling, Jane Hung, Rex Wang, Callum Tromans-Coia, Bin Li, Edward L Evans, Kevin W Eliceiri, Peter Horvath, Anne E Carpenter, Beth A Cimini

## Abstract

Deep learning has greatly accelerated research in biological image analysis yet it often requires programming skills and specialized tool installation. Here we present Piximi, a modern, no-programming image analysis tool leveraging deep learning. Implemented as a web application at Piximi.app, Piximi requires no installation and can be accessed by any modern web browser. Its client-only architecture preserves the security of researcher data by running all computation locally. Piximi offers four core modules: a deep learning classifier, an image annotator, measurement modules, and pre-trained deep learning segmentation modules. Piximi is interoperable with existing tools and workflows by supporting import and export of common data and model formats. The intuitive researcher interface and easy access to Piximi allows biological researchers to obtain insights into images within just a few minutes. Piximi aims to bring deep learning-powered image analysis to a broader community by eliminating barriers to entry.

## Introduction

Image analysis has played a crucial role throughout the history of biological and medical research. Within the last few decades, the amount and variety of image data generated and analyzed has increased dramatically, requiring powerful tools to automate tasks such as cell segmentation or classification. These tools are essential in the automated analysis of image data but also allow researchers to focus on more sophisticated tasks than manually annotating or classifying cells. Currently, the bioimaging community relies heavily on popular Graphical User Interface (GUI)-based open-source applications relying on classical image processing and machine learning, such as general-purpose tools like ImageJ ^1^ / Fiji ^2^, CellProfiler ^3^, Icy ^4^, and napari ^5^ and more specialized tools such as ilastik ^6^, CellProfiler Analyst (CPA) ^7^ and Advanced Cell Classifier (ACC) ^8^. Despite the wide adoption of various tools supporting image analysis in biomedical research, these programs all come with their limitations. Most of them require the installation of a local program or setting up a controlled computing environment which not only adds an additional hurdle for many researchers but also often limits the available platforms. Additionally, many tools are complicated to use, lack sufficient researcher documentation, have slightly outdated researcher interfaces, or are designed for specific use cases only.

The last decade of image analysis has seen a wide adoption of deep learning-based tools and algorithms, further accelerating bioimage analysis ^9^. Cellpose ^10^ is a highly popular stand-alone deep learning segmentation tool for bioimage analysis and offers powerful pre-trained cell segmentation models, and significant efforts have been made to deploy deep learning tools through napari; Fiji (by the DeepImageJ project ^11^); CellProfiler ^12^; and in ilastik, QuPath ^13^ and other tools via the Bioimage Model Zoo ^14^. However, the fact remains that using most deep learning workflows requires comfort in writing code and running it through a command line interface or in complex deployment or programming environments which can require additional steps to troubleshoot dependency installation ^15^. The required programming skills limit the researcher base and thus the potential applications of deep learning in bioimage analysis ^16^. These practical limitations generally limit modern applications of deep learning for less-computationally-comfortable researchers in many fields.

Here we present Piximi, an open-source, no-code, and modern web-based deep learning application. Our goal was to offer a complete deep learning bioimage analysis workflow in a single, easy-to-use tool.

In this initial version, Piximi offers four core functionalities:

1. Classifier: allowing researchers to easily label images or objects within images, such as cells, and train a classifier to recognize them.
2. Annotator: providing multiple flexible tools for creating object bounding boxes and segmentation outlines to allow researchers to annotate images directly in Piximi for use in downstream tasks.
3. Segmenter: allowing researchers to find object within images using pre-trained instance segmentation and object detection networks
4. Measurements: capturing various properties of images or entities identified therein, such as size, shape, and intensity. Measurements can then be plotted to examine distributions or between-group differences.

Piximi is implemented as a freely available web application (https://www.piximi.app/). Thus, unlike many other tools in the field of image analysis, it is available on any operating system (PC, MAC, Linux, and mobile) through any modern browser and without any installation required. In addition to its ease of access, we aimed to create an easy-to-use, no-code application that democratizes science by allowing everyone to leverage deep learning on their projects without any programming skills.

Piximi offers multiple tools in a single application, which allows researchers to complete full workflows within a single tool rather than switching between different programs for individual sub-tasks. It relies on commonly used data formats such as the COCO object detection format, to guarantee interoperability with existing tools and frameworks. Generally all computation is done on the local device and no image data is sent to a server (client-only); certain models will run data on a remote server but only with researcher permission. This privacy-preserving implementation is a key feature allowing use on sensitive or confidential data such as medical images, or in low-resource settings where internet access is occasionally available but not consistently reliable.

Piximi can also be used across a variety of image domains and was intentionally designed to support a large variety of use cases. Early adopters have explored this tool for various imaging tasks including cell type classification, annotation of radiology images, and application of segmentation models on histology images. By enabling deep-learning-based image analysis for a range of researchers who were previously excluded due to a lack of computational skills, allowing them to turn images into answers for a variety of scientific questions.

## Results

### Classifier

The classifier module offers an intuitive and easy way of classifying large numbers of images. Unlike related tools such as CPA or ACC, Piximi does not require a separate feature extraction step but instead works on the raw images. Researchers can import images into Piximi from a local directory, and label a subset of them (that is, point out to the software which images correspond to which categories of interest). These labels are used as ground truth in the training process, so that the deep-learning classifier learns to predict labels for unknown images. Hyperparameters of the model can be adjusted, such as loss function, learning rate, number of epochs, and the train and validation split among the labeled images can be defined.

### Simple classification example project

To demonstrate the ease of use of classifying images using Piximi we created an example project based on a set of HeLa cell images showing the differential localization of the healthy versus disease-associated variant of the PLP1 protein ^17^.

To classify unlabeled images as either reference (healthy) or variant (disease-associated), we used Piximi to train a classifier. Using a training set of only 50 labeled cell images each, the classifier correctly identifies the majority of images, reaching an F1 score of 0.81 after just 50 epochs of training. The training process on Piximi in the browser finishes in only a few minutes on a typical laptop.

**Figure 1:**
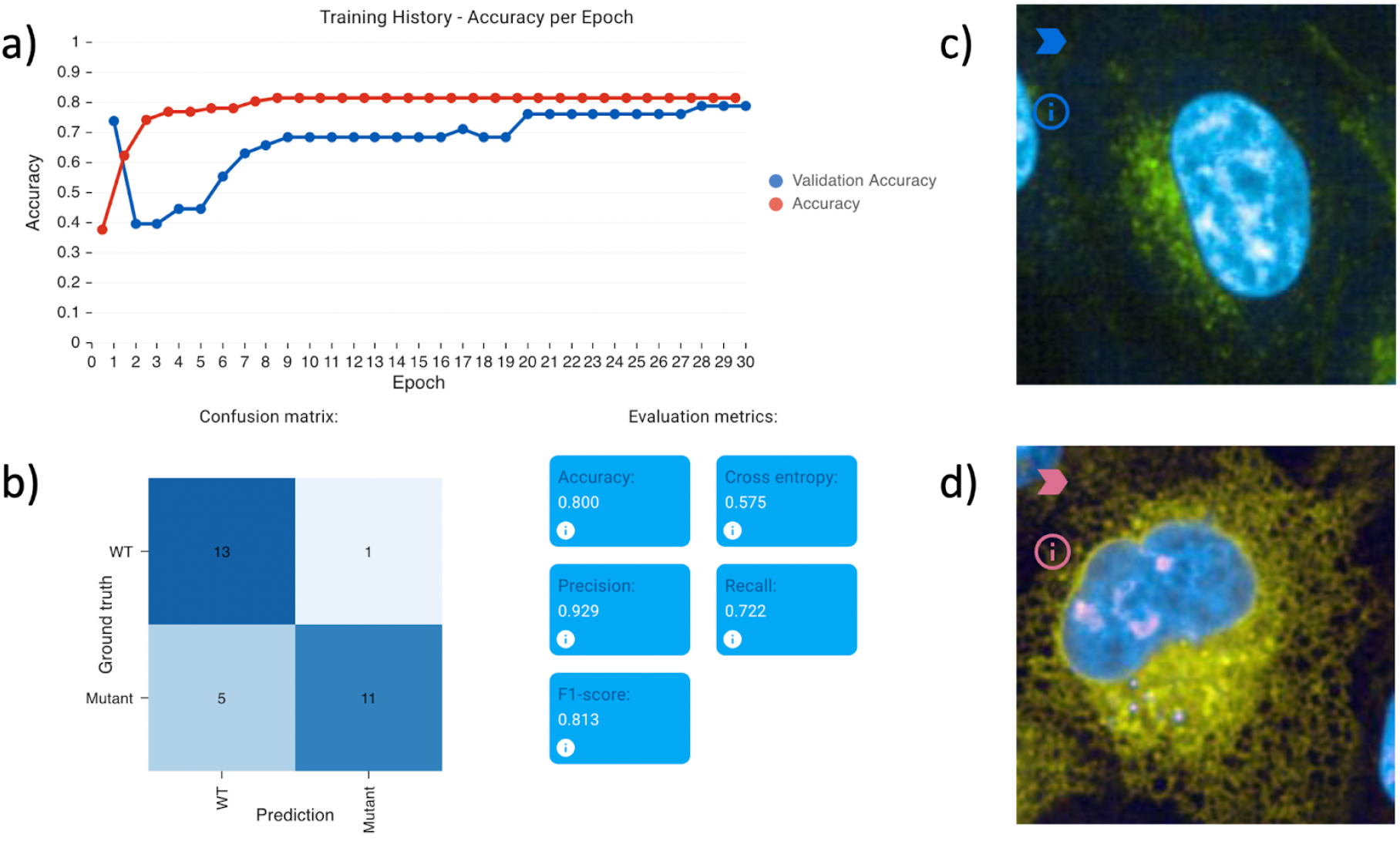
Evaluation results on the PLP1 localization classification example project, provided within Piximi (Open -> Project -> Example Project -> Human PLP1 localization).Two channels were imaged during the experiment, channel 0 shows the targeted PLP1 protein (green), and DNA (blue) is imaged in channel 1. The dataset contains 129 images labeled as either wild-type or mutant. a) Shows the training history and b) shows useful evaluation metrics. c) and d) show two examples of the prediction result for reference and variant respectively.

### Continuous training/human-in-the loop

Piximi supports continuous training of models. This enables training models more efficiently by applying a “human-in-the-loop” approach: after annotating a small subset of images, a researcher can specify the classifier and train it, then show the predictions of the network. Classification errors can be corrected by reassigning the respective images to the correct label. Restarting the training procedure resumes training where it previously stopped, using the now-expanded training set.

This iterative process can be repeated until the evaluation results are satisfactory. This approach typically results in better overall results while requiring considerably less manual annotation of images ^10,18,19^.

**Figure 2:**
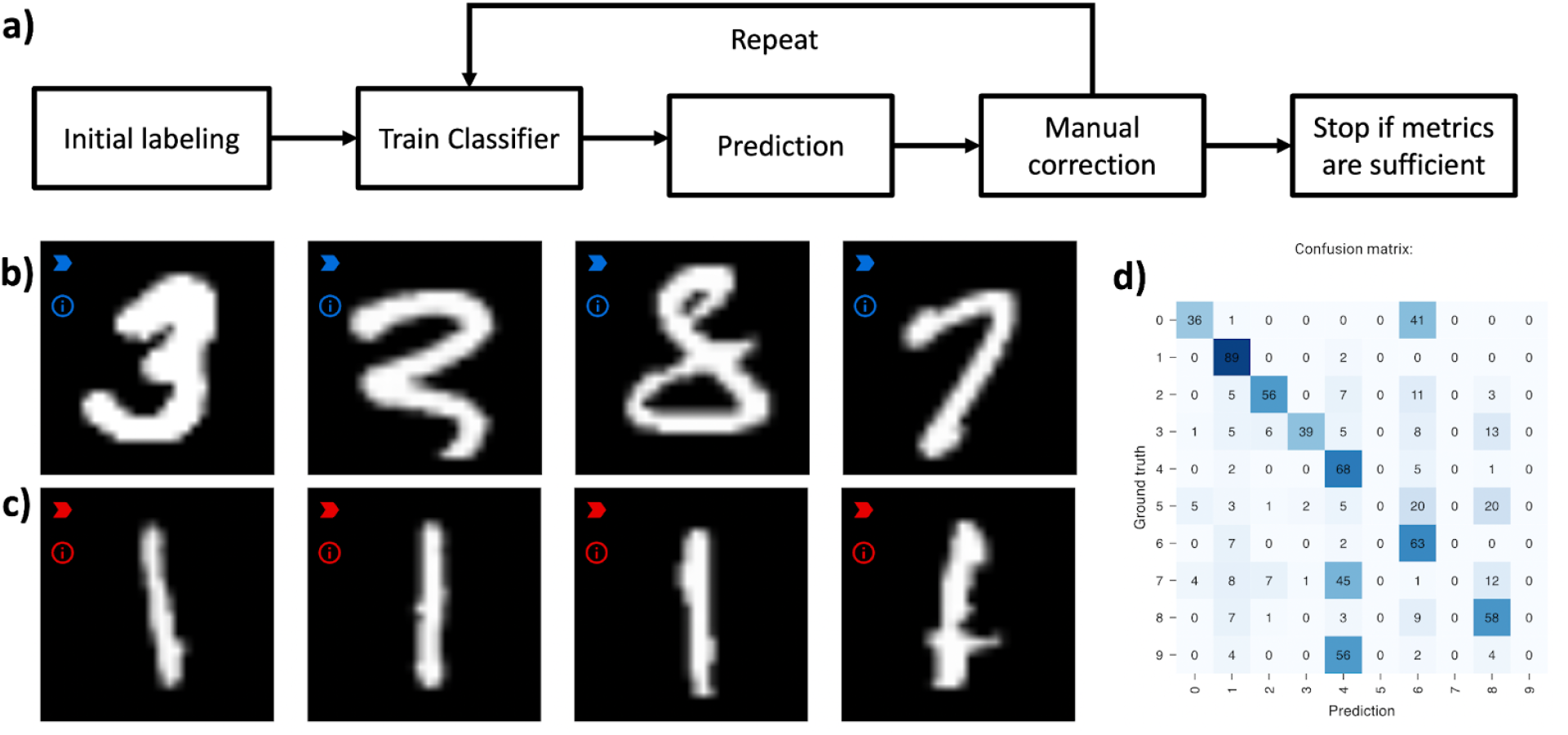
Example workflow for human-in-the-loop training using MNIST ^20^. a) Abstraction of steps in a human-in-the-loop learning procedure. b) Example MNIST digits predicted as “3” after a few iterations. The misclassified digits can be corrected by assigning them to the right label (“8” and “7”). c) Example of correctly predicted digits. Images with easy patterns such as “1”s are mostly correctly identified after only a small number of manually labeled examples. d) Researchers can inspect the confusion matrix available in Piximi to see which classes are misclassified the most, to focus their efforts where most needed.

### Annotator

A common step in most bioimage analysis workflows is the boundary annotation of objects in images, such as cells. Recent years have seen a surge in useful annotation tools such as QuPath ^13^, ITK-SNAP ^21^, and various napari plugins.

Object annotations are especially important for follow-up deep-learning tasks such as segmentation or cell-type classification which first require the identification of individual cells or nuclei.

We implemented an annotator as part of Piximi which follows the designed principles outlined in the introduction. Without any installation steps, Piximi offers a selection of tools to annotate objects of interest. Simple annotation tools such as rectangular, elliptical, polygonal tools allow the user to manually draw simple geometric shapes around the objects of interest, while more sophisticated tools automatically mark the region of interest. The quick-selection tools use the SLIC algorithm ^22^ to calculate so-called superpixels in the images, which typically encompass a single object. Similarly, the “Color tool” uses a flood fill algorithm to directly identify objects based on researcher-selected thresholds. This tool is particularly useful for objects visually identified by color intensities, such as stained compartments of cells or hyper-intense areas within the image. For grayscale images in particular, it might often be enough to identify objects using a simple threshold. For such cases, the Piximi annotator offers a threshold tool that annotates regions below the specified threshold within the bounding box. Annotation can easily be corrected by manually adding or subtracting regions using any available tool.

The annotator module also serves as an image viewer for inspecting images more closely. Piximi supports both multichannel as well as 3D images. Researchers can slide through the z-dimension of the images and individually annotate each slice. Researchers can select which channels to show or hide and assign colors to specific channels.

In keeping with the core design principle of interoperability, annotations in Piximi can easily be exported and used in subsequent machine learning pipelines, for example to train deep learning models using common frameworks such as TensorFlow ^23^ or PyTorch ^24^. Available commonly used data formats include labeled instance masks, binary instance masks, binary semantic masks, and COCO-formatted ^25^ annotation data.

**Figure 3:**
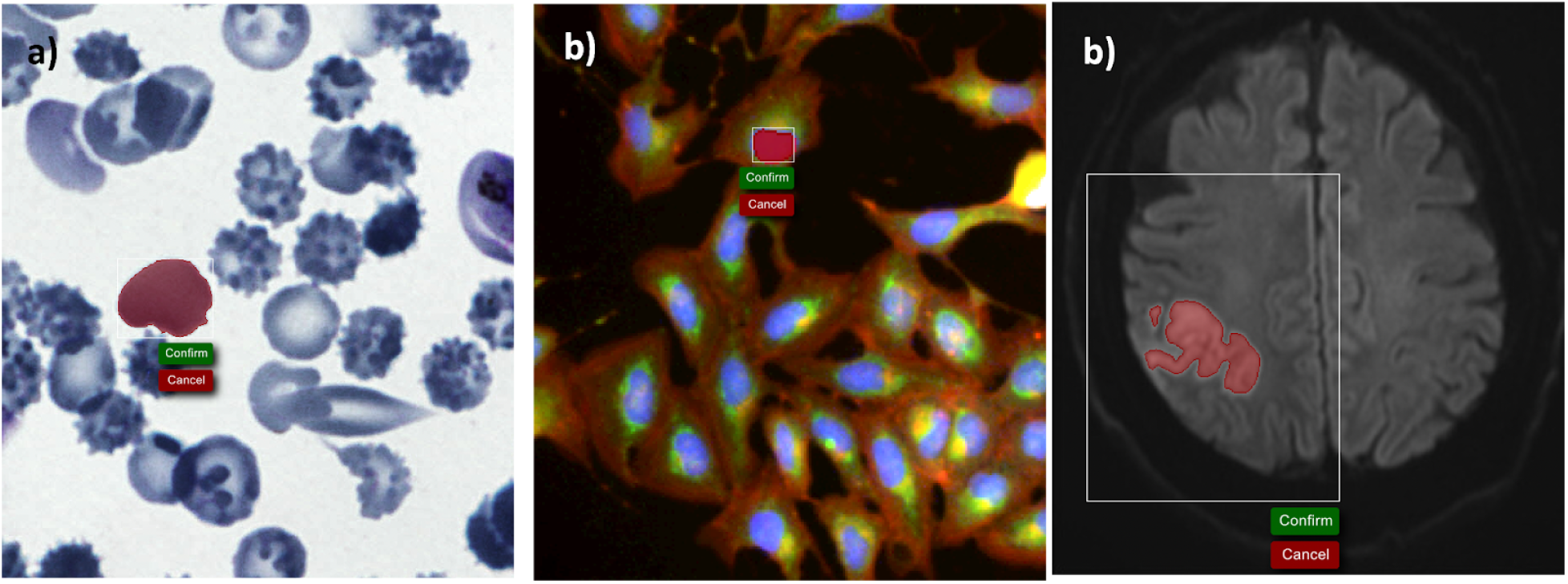
Use of the Piximi annotator on brightfield (left), fluorescence (center) and MRI (right) data.

### Segmenter

Segmentation is another common task in bioimage analysis. Many downstream analysis tasks including cell type prediction or feature extraction require the segmentation of individual cells within the tissue or cell population. Options include well-known segmentation algorithms such as distance-based watershed, complex segmentation tools such as CellProfiler ^3^, and many deep learning-based segmentation algorithms including Cellpose ^10^ or StarDist ^8,26^. Deep learning algorithms regularly outperform classic image analysis pipelines ^27^. However, deep learning pipelines are often complex to set up and require the installation of various dependent packages.

The segmentation module within Piximi offers a selection of pre-trained segmentation models that allow researchers to quickly identify nuclei or cells. A pre-trained model is selected from the available model zoo and inference is applied to the images opened in Piximi. The web-based implementation and easy-to-use interface allow the obtaining of segmentation results of cells within just a few minutes. The ease of use of these models in Piximi is especially notable for small datasets where setting up a custom pipeline would be infeasible or overly onerous.

Currently, the annotator offers five pre-trained segmentation models. There are two segmenters which work on hematoxylin and eosin (H&E) stained images: StarDistVHE to identify nuclei in hematoxylin and eosin (H&E) stained images, as well as a compact UNet ^28^ which segments intestinal glands trained on the Gland Segmentation in Colon Histology Images Challenge Contest (GlaS) ^29^. To showcase non-biological applications as well as the ability to use multi-class object detectors, we also include COCO-ssd, which identifies objects in “natural images” (or photographs) of 80 different classes (such as humans and kites) using the COCO format. We also support two segmenters for fluorescence microscopy; StarDist, and Cellpose. Cellpose is currently unique in that it runs on the AI4Life project’s BioEngine ^30^ server while StarDist, like other Piximi models, runs client-only in the user’s own browser without data leaving their machine. An alert is shown when selecting Cellpose which warns users that images will leave their machine. While Piximi will remain a primarily client-only program, supporting an optional remote server for certain specific models allows Piximi to run highly memory-intensive models or models that contain elements which are cumbersome to port to Tensorflow.Js ^31^,such as models containing custom postprocessing Python libraries.

Segmentation results are accessible as Piximi object annotations, allowing researchers to edit and/or export segmentation masks as can be done with manually created annotations. While Piximi does support multiclass object finding (in which a researcher could simultaneously separately identify different object classes such as mitotic cells and non-mitotic cells), the object annotations are also available in the classifier tool as well, allowing a researcher to run an initial segmentation model that grabs all objects (such as cells) and then separate them into classes (mitotic and non-mitotic).

**Figure 4:**
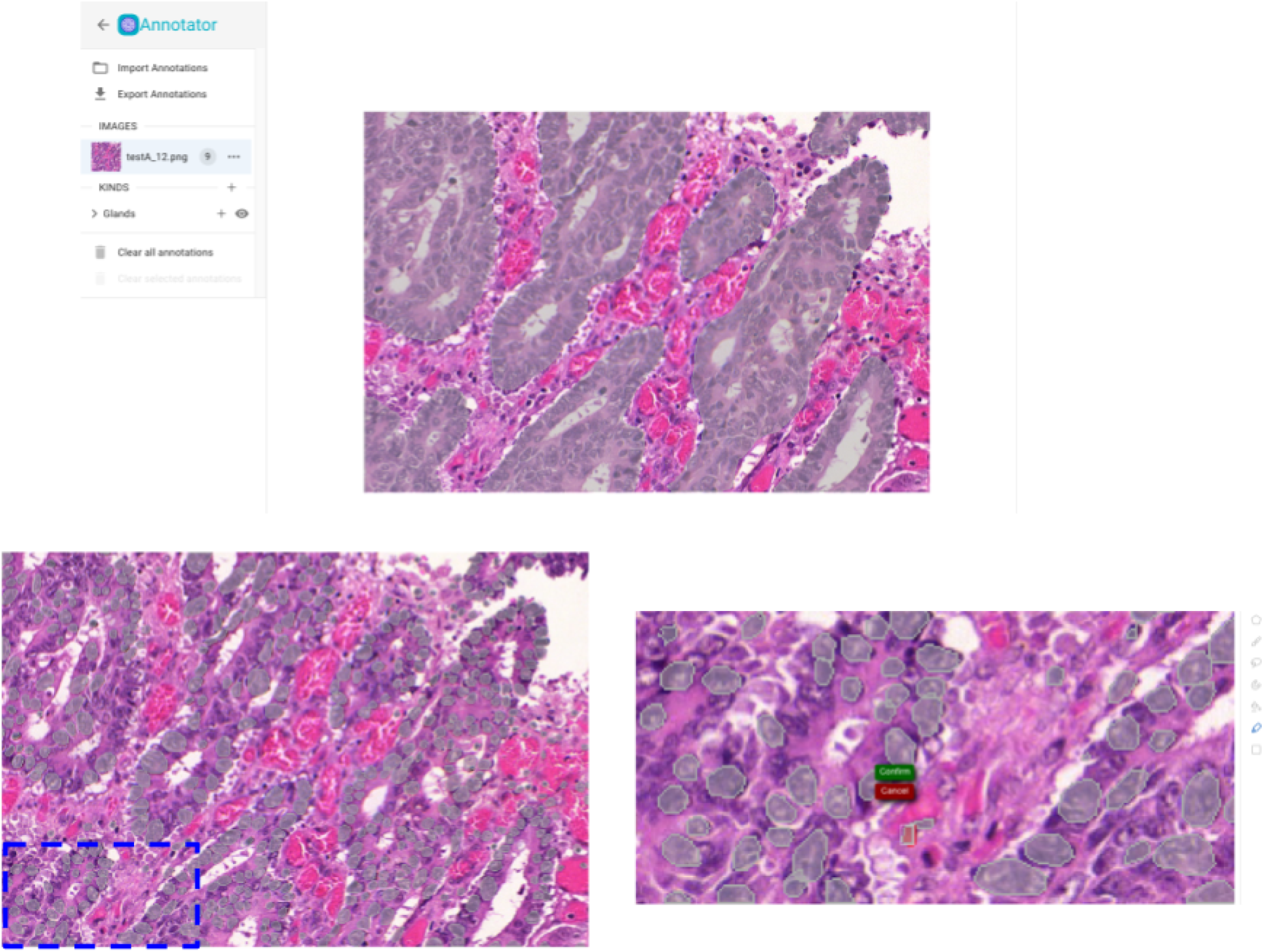
Segmentation results on a test set image from the Gland Segmentation in Colon Histology Images Challenge Contest. Top: results of the Gland segmentation model. Bottom: results of the StarDistVHE nuclear detection model. Left: unedited segmentation results. Right: missing segmentations can be easily added using Piximi’s quick-selection tool.

### Measurements

To offer researchers a comprehensive end-to-end experience in understanding the experimental results within a single application, Piximi features a measurement module that provides insight into project images and annotation objects by quantifying key information and attributes. Users perform measurements by first selecting a group of images or objects to be measured, referred to as a “split”. Currently users can specify splits by first selecting the object kind they’re measuring (such as “image”, “cells” or “glands”), then they further narrow the selection by specifying the category the object or images belong to, and/or the training partition. For instance, a split may be all of the “cell membrane” objects categorized as “LargeDiffuse”, or all of the imported images used for validation in a model. The measurements available are split into two groups, image level measurements and object level measurements. The image level measurements include measurements of the intensity of the image (total, median, standard deviation, etc.) and the object level measurements include measurements of object geometry (area, perimeter, sphericity, etc.). See the Methods for the full list of measurements and their implementations.

**Figure 5:**
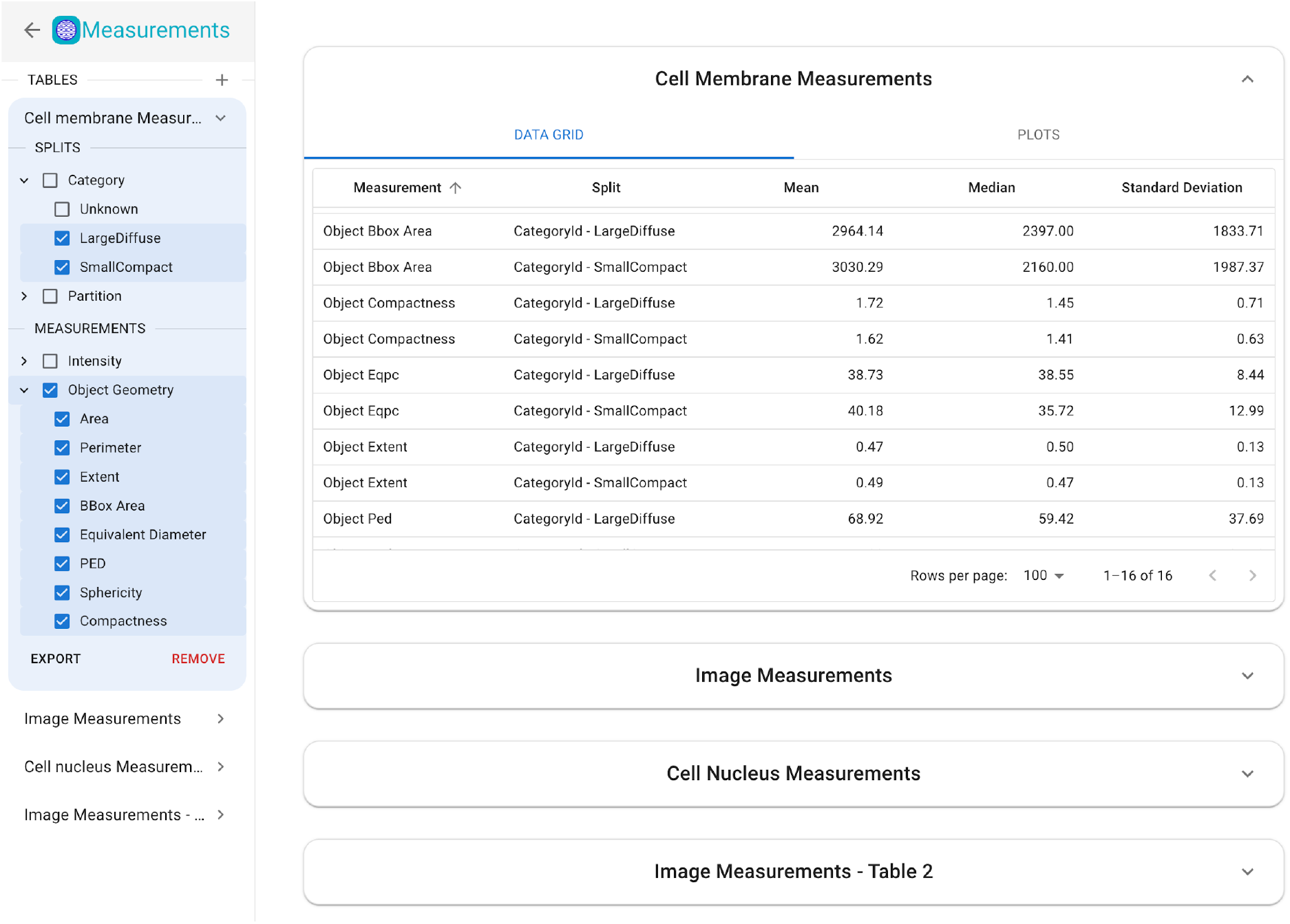
In-app web measurement functionality in Piximi. Image and object measurements can be made and exported; objects or images which have been assigned to categories will show the average, standard deviation, and median value within each category.

Selecting the splits and measurements will populate the table with the total, median, and standard deviation statistics for the measurements for each split. Users can sort and filter the rows of the table using the functions provided in the column headers. Users can create multiple tables at a time and easily switch between them and collapse the ones they aren’t working in.

Following the design principle of interoperability, measurements can be exported to a CSV file to use for further downstream analysis. Exported measurements can easily be used in other data exploration tools like Morpheus ^32^ or CellProfiler Analyst ^7^. The CSV file contains the per image/object measurements.

In addition to the numerical table, users also have the options to visualize their measurements using a selection of plots – histogram, swarm, and scatter – and apply color and size mapping. Users can specify the parameters for the plots in the left section of the plot tab. Each plot has a associated group of parameters, some which vary depending on the plot: the parameters for histogram (Figure 6A) are the x-axis measurement and the number of bins, the scatterplot (Figure 6B) takes x- and y-axis measurements, as well as an option to map the mark size to a measurement and color to a split, and the swarm plot (Figure 6C) takes a y-axis measurement, swarm group which is associated with a split (category or partition), mark size, and an option to toggle a measurement statistics overlay which displays the median, std, and upper and lower quartiles of a group (Figure 6D).

**Figure 6:**
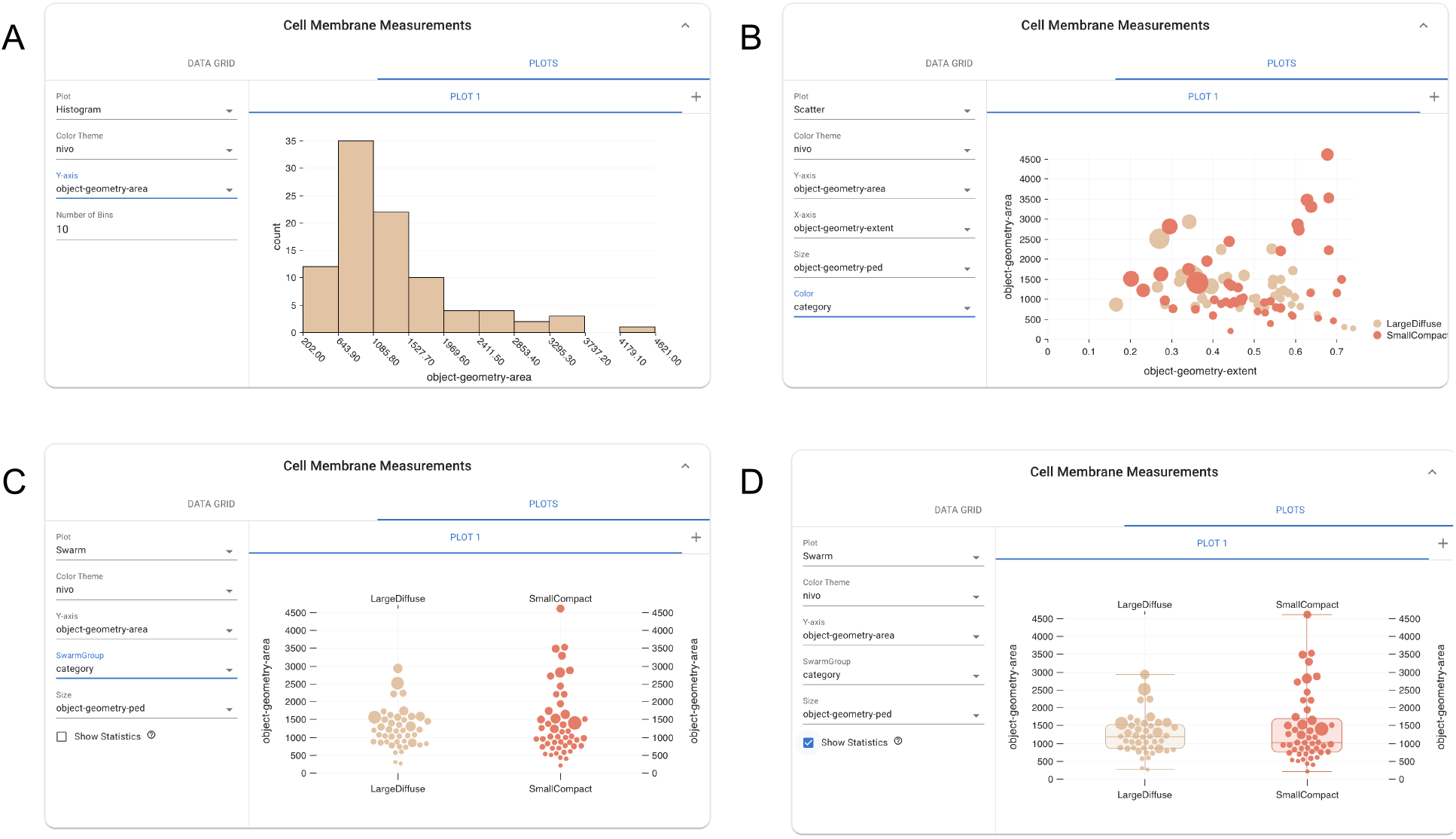
Snapshot of the histogram (A), scatterplot(B), swarm plot(C), and the swarm plot with a statistics overlay(D) as well as the associated parameters for each of the plots.

Users can select from a large number of color themes to apply to the plots, and each plot will have its own associated theme. It is also possible to create multiple plots at the same time. Each new plot opens in a new tab within the table, and each tab can be renamed to allow for greater clarity when working with multiple plots. Plots which are no longer needed can be easily deleted using the delete button in the tab. Users will also have the ability to save the plots then generate using Piximi.

### Data and model interoperability

Piximi was developed with interoperability with other tools in mind. Image labels, predictions, and measurements can easily be exported and used for further downstream analysis using various tools.

The architecture graph and weights of any models trained in Piximi can also be downloaded and imported into other tools and workflows. This enables training an initial classifier on Piximi using the human-in-the-loop approach to limit manual annotation of images, then exporting the model to run inference on large datasets on a cluster or in the cloud (which are not currently suitable for processing by Piximi). Conversely, models developed and trained in a Jupyter Notebook or other Python environment can also be imported into Piximi. For instance, we trained a UNet for intestinal gland segmentation in H&E stained images of colorectal tissues. This trained model was then imported into Piximi, enabling the segmentation workflow to be run directly in the Piximi browser application, circumventing the need for a complex Python environment or a computing server. This enhances model accessibility for non-computational researchers as well as substantially increases reproducibility.

## Discussion

Here we described Piximi, a modern web application that offers an easy-to-use, intuitive non-code interface to perform common image analysis tasks. In this first version, Piximi offers four core modules: a deep learning classifier, an image annotator, pre-trained segmentation models, and measurements. These modules are not implemented as standalone tools but are interoperable within Piximi. All analysis results and data, including classification prediction, labels, measurements, and annotation or segmentation masks can easily be exported to common data formats, yielding a reproducible record of analysis and allowing results to be used with other tools and workflows. This interoperability with the broader image analysis community is a key feature of Piximi.

Piximi is implemented as a client-only web application without any server component. Unless the user chooses a pretrained model only supported remotely, no images or data are uploaded to a server and all computation is done locally. In addition, Piximi can be served via a Docker container which enables institutional researchers to deploy Piximi within their local network easily.

The implementation of Piximi as a web application also eliminates the need for an installation process. Additionally, Piximi can be accessed from any modern web browser and, beyond accessing example data sets or remote server tools like Cellpose, continuous internet connection is not needed following initial loading.

However, implementing Piximi as a web application creates certain limitations. Most notably the performance of training deep learning models in the browser is limited compared to non-web GPU workflows. Large datasets or large images might be slower to handle compared to comparable traditional applications. In general, web applications like Piximi also currently lack access to the local file system and direct interaction with image analysis libraries written in Java or Python. Instead, most image analysis features in Piximi are re-implementations in JavaScript. Nevertheless, major developments in web applications such as File System Access API or WebAssembly (WASM) will benefit applications like Piximi.

Future work on Piximi will focus on both expanding the feature set and improving the performance and researcher experience of Piximi. Performance improvements will include support for large images and larger overall datasets. In addition to pre-trained segmentation models, future versions of Piximi will also support trainable segmentation models. We also aim to include more efficient human-in-the-loop training by providing a researcher interface that guides through this process—by identifying hard-to-classify images and guiding the researchers through the process, the training process will converge faster and likely produce better results.

In conclusion, we presented a new, modern web application for image analysis. We showed particular use cases of Piximi for quickly and efficiently running cell type classification, nuclear segmentation, and tissue segmentation using the included deep learning tools.

Although initially created to bring bioimage analysis tools to a broader audience, Piximi can be used for any other research field as well. Its web-based implementation and intuitive no-code researcher interface enable us to bring deep learning-based image analysis tools to researchers who were previously excluded or faced major barriers in accessing such networks. We believe Piximi’s powerful features, ease of access, and ease of use help it fill a unique niche in the image analysis ecosystem.

## Online Methods

### Implementation details

Piximi’s source code is publicly available on GitHub and is provided under a BSD-style open-source license.

Piximi is implemented using state-of-the-art web development tools. The application is written in TypeScript using the React library for the UI. Libraries such as React allow us to develop custom components which streamlines development as well as providing performance benefits which stem from the ability to monitor and cut down on re-renders. In order to accomplish the goal of providing an intuitive user experience with a low barrier to entry, styling is implemented via the MaterialUI library, which adheres to the widely used open-source Material Design system developed by Google.

Application state and data are managed using the Redux Library, which allows for the creation of a “store” which holds the data and provides a simple api for Create, Read, Update, and Delete (CRUD) functionality. Since this store is accessed by many different components used in the application, Redux provides methods and guidelines for ensuring only deliberate mutation of the store. The otherwise immutability of the store guarantees that data and state stay consistent and in-sync between app re-renders and the various components. Application data (images, annotations, categories, and images/objects (collectively referred to as “kinds”)) are kept in the store within modified “entity adapters”. The entity adapter is provided by Redux as a way to quickly access data by storing it in id-based lookup tables along with providing a list of the ids within each adapter. We modified the entity adapters to allow for updating the data in a non-permanent way, storing only the changes made until the user decides to save those changes or revert them.

Using TensorFlow.js ^31^, we are able to store arbitrary numerical arrays as tensors. Therefore, in addition to the training and inference of models, we are able to store all image data in much more compute-efficient data structures. This provides a distinct advantage over using JavaScript’s primitive arrays as it allows us to carry out image processing algorithms with gpu-bound, vectorized operations at the whole-array level, rather than needing to sequentially iterate over each individual element in a multidimensional array and perform cpu-bound arithmetic and logical operations on them.

The use of TensorFlow.js also allows for the selection of multiple compute backends. Currently, for the browser, this includes cpu-bound operations via the “cpu” (vanilla javascript) and “wasm” (XNNPACK library compiled to WASM) backends and a “gpu” backend with gpu-bound operations which are implemented in OpenGL Shading Language (GLSL) and routed through WebGL. There is an in-progress “webgpu” backend for gpu-bound operations which utilizes the more modern and performant WebGPU API available in most modern browsers.

Since javascript is single-threaded, computationally intensive tasks run the risk of freezing up the UI, preventing users from further interaction with the application until the task is complete. We therefore employ Web Workers in various points in the code, which allows Piximi to run scripts in background threads without affecting the main UI. Data is sent back and forth between the main thread and the worker using a system of messages, allowing Piximi to provide useful information about the running process during calculation. An added benefit of using Web Workers is the ability to spawn a process in Piximi and switch to another tab in the browser while the computation takes place, increasing the efficiency of a user’s workflow.

### Measurement implementation

Measurements are calculated using a mix of tensorflow and custom functions, and are computed within web workers to keep the main thread unburdened. As mentioned previously, measurements are separated into two groups, image level measurements and object level.

The image level encapsulates several measurements for the image intensity and are calculated for each channel of the images. These measurements include: total, mean, median, standard deviation, mean absolute deviation (MAD), lower quartile, and upper quartile of the pixel intensities. The MAD is computed as

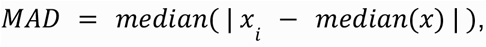

and the lower(upper) quartile is defined as the intensity value of the pixel for which 25%(75%) of the pixels in the object have lower values.

Unlike image level measurements, which can be performed on both images and objects, object level measurements require that the object have a mask generated by Piximi, and thus cannot be performed on pure images uploaded to Piximi. The object measurements currently implemented are used to measure geometrical features of the objects:

- Area: sum of the number of pixels in the object
- Perimeter: total number of pixels around the boundary of the object
- Bounding Box Area: total number of pixels in the bounding box
- Extent: the proportion of the pixels in the bounding box that are also in the region, computed as

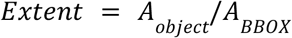
- Equivalent Diameter: the diameter of a perfect circle with an area equal to the area of the object

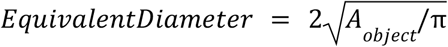
- Diameter of equal perimeter (PED): the diameter of a circle whose perimeter equal to that of the object

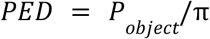
- Sphericity: measures the ratio of the perimeter of the equivalent circle, *P*_*ECPC*_, to the real perimeter of the object, *P*_*real*_. The result ranges from 0 (irregularly shaped) to 1 (spherical).

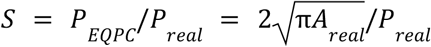
- Compactness: the degree to which the object is compact, with circular shapes being most compact and irregular objects will have a value greater than 1, increasing in irregularity. Measured as the inverse of sphericity.

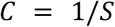

All the measurements are stored per image/object within Piximi, and only need to be recomputed if the underlying data changes.

### Model Zoo

Piximi’s classifier offers two different classification model types: “SimpleCNN” is a simple implementation of a fully convolutional network. Additionally, researchers can use a Mobilenet, ^33^ a small and efficient deep-learning model for object classification. These networks were specifically developed to achieve high classification accuracy while having significantly fewer parameters and lower inference latency than comparable models. Mobilenets are often used in mobile applications and web services. Piximi uses a frozen pre-trained mobilenet and trains an additional layer mapping from the last hidden layer to the specified output shape.

The first version uses the four pre-trained segmentation models mentioned above: COCO-SSD, StarDistVHE, and a UNet model for identifying glands in tissue images.

### Data Formats

Piximi supports common image files such as png, jpeg and Tiff. Tiff files can have an arbitrary number of z-dimensions or channels. On importing the images, the researcher specifies the number of channels for hyperstack images. Additionally, Piximi supports uncompressed DICOM files as well.

Projects on Piximi, including labels and images, can be downloaded into a Zarr file to save projects on local disk. Annotations and segmentation masks can be exported using common formats including COCO formatted JSON or binary schematic masks.

Models are stored in one of two TensorFlow.js formats, Layers Models or Graph Models, in the form of a json file containing the model description (layers, ops, inputs, outputs) and sharded binary files containing model weights. Layers Models are fully trainable, whereas Graph Models are optimized for inference only. Both formats can be converted to using the tfjs-converter CLI tool utilizing one of several input formats, such as TensorFlow SavedModel, Keras HDF5 or Keras SavedModel. Likewise the Layers Model format can be converted back into Keras HDF5 or Keras SavedModel formats. The tf2onnx tool also allows us to convert TensorFlow.js models to an ONNX capable format. Likewise, although more complicated, ONNX may be utilized to convert Torch (.pt/pth) formats to a suitable Keras format, and then ultimately to a TensorFlow.js Graph model.

## Acknowledgements

The authors also want to acknowledge Pearl Ryder, Frances Hubis, and all the participants of the CytoAI and Piximi hackathons. Members of the Horvath, Carpenter-Singh, and Cimini labs are also gratefully acknowledged for their contributions to the paper, the code, the UI/UX, and the documentation.

## Funding

The work was supported by grants TKP2021-EGA09, Horizon-BIALYMPH, Horizon-SYMMETRY, Horizon-SWEEPICS, H2020-Fair-CHARM, CZI DVP, HAS-NAP3,OTKA-SNN no. 139455/ARRS to PH, National Institute of General Medical Sciences P41 GM135019 to BAC, AEC, and KWE, R35 GM122547 to AEC, grant numbers 2020-225720, 2018-183451, and 2021-238657 to BAC from the Chan Zuckerberg Initiative DAF, an advised fund of Silicon Valley Community Foundation. The funders had no role in study design, data collection and analysis, decision to publish, or preparation of the manuscript.

## Notes

https://www.piximi.app

### Competing Interest Statement

The authors have declared no competing interest.

### Summary of Updates

The manuscript now reflects additional updates to Piximi's measurement capabilities, including graphing

